# Cerebellum involvement in visuo-vestibular interaction for the perception of gravitational direction: a repetitive transcranial magnetic stimulation study

**DOI:** 10.1101/2025.03.09.642248

**Authors:** Keisuke Tani, Hiroaki Tanaka, Akimasa Hirata, Nobuhiko Mori, Koichi Hosomi, Akiyoshi Matsugi

## Abstract

Accurate perception of the direction of gravity relies on the integration of multisensory information, particularly from the visual and vestibular systems, within the brain. Although a recent study of patients with cerebellar degeneration suggested a cerebellar role in visuo-vestibular interaction in the perception of gravitational direction, direct evidence remains limited. To address this gap, we conducted two experiments with 42 healthy participants to evaluate the impact of repetitive 1-Hz transcranial magnetic stimulation (rTMS) over the posterior cerebellum on visual dependency, quantified by the subjective visual vertical bias induced by rotating optokinetic stimulation (OKS). Electric field simulations in high-resolution head models were used to ensure focal stimulation of the cerebellum at the group level. The results demonstrated that repetitive transcranial magnetic stimulation (rTMS) applied to the cerebellar vermis significantly attenuated the OKS-induced shift in visual vertical (SVV) bias. This effect was not observed when stimulation was applied to the early visual cortex (V1–2) or the cerebellar hemisphere. Also, the vermis rTMS had no effect on the judgement precision in the absence of visual motion cues, suggesting that the rTMS may reduce visual weight in visuo-vestibular processing by increasing visual motion noise rather than affecting vestibular function. These findings suggest a direct involvement of the cerebellar vermis in the visuo-vestibular interaction underlying the perception of gravitational direction, providing new insights into cerebellar contributions in human spatial orientation.

**Significance Statement:** The cerebellum has been implicated in multisensory integration for spatial orientation, but its direct role in visuo-vestibular interactions remains limited. Using 1-Hz rTMS, we demonstrated that stimulation of the cerebellar vermis significantly reduced visual dependency in the perception of gravitational direction, as measured by the subjective visual vertical bias induced by rotating optokinetic stimulation. This effect was absent when adjacent areas, such as early visual cortex and the cerebellar hemisphere, were stimulated. These results suggest that the cerebellar vermis is directly involved in visuo-vestibular interaction, providing new insights into the cerebellar contribution to spatial orientation in humans.

## Introduction

Awareness of the direction of gravity provides a fundamental basis for the perception of external events and motor control on Earth. A large body of psychophysical evidence has shown that the brain integrates different types of sensory information, especially vestibular and visual information, to internally estimate the direction of gravity, likely in a statistically optimal manner (Niehof et al., 2019; Angelaki et al., 2009; see also Kheradmand & Winnik, 2017 for a review). However, the neural substrates underlying visuo-vestibular interaction in the context of the perception of gravitational direction are largely unclear.

The otolith organs in the inner ear, comprising the utricle and saccule, respond to linear acceleration and constitute an essential sensor for detecting gravitational direction with respect to the head (Goldberg et al., 2012). However, because gravitational (tilt) and translational accelerations are physically indistinguishable (Einstein, 1907), the otolith signal provides ambiguous information regarding the gravitational direction. To resolve this (tilt–translation) ambiguity, the brain seems to track the direction of head tilt relative to gravity on the basis of angular velocity signals from the visual system, as well as the semicircular canal, and integrate this information with otolith signals (Laurens & Angelaki, 2011; Dakin et al., 2018; MacNeilage & Glasauer, 2018; Diaz-Artiles & Karmali, 2021). In fact, it is known that large-field angular visual motion (i.e., rotating optic flow) leads to a strong bias in the perceived direction of gravity towards its direction (Dichgans et al., 1972).

Recently, Dakin et al. (2018) showed that the bias of the perceived direction of gravity induced by large-field visual motion was much greater (interpreted as “increased *visual dependency”*) in patients with spinocerebellar ataxia type 6 (SCA6) and slowly progressive degeneration limited to the cerebellum than in age-matched healthy participants. This finding suggests that the cerebellum subserves the visuo-vestibular interaction for the perception of gravitational direction. However, recent studies have reported reduced functional activity not only in the cerebellum but also in various cortical regions (Kang et al., 2017), as well as changes in functional/effective connectivity between the cerebellum and cortical regions as symptoms progress (Kang et al., 2017; Falcon et al., 2015). These findings suggest that the increased visual dependency observed in SCA6 patients may be attributed to compensatory functional changes in cerebral cortical areas, such as the visual cortex, following cerebellar degeneration. Therefore, whether the cerebellum is directly involved in visuo-vestibular interaction remains unclear.

To address this issue, we performed two experiments in healthy participants using low-frequency (1-Hz) repetitive transcranial magnetic stimulation (rTMS), which can induce focal and short-term modulation of neural activity in targeted brain regions (Pascual-Leone, 1999; Kvamme et al., 2022). Neurophysiological studies have demonstrated that cerebellar brain inhibition, which reflects the inhibitory mechanism of the cerebellum on the motor cortex, is suppressed (i.e., de-inhibited) by 1-Hz rTMS over the cerebellum (Popa et al., 2010; Fierro et al., 2007; Oliveri et al., 2005). Additionally, 1-Hz cerebellar rTMS has been shown to disrupt motor learning (Lien et al., 2022; Matsugi et al., 2023) and reduce perceptual and cognitive performance (Fierro et al., 2007; Schutter et al., 2009). Therefore, 1-Hz rTMS over the cerebellum seems to generally suppress cerebellar function (Gatti et al., 2023).

Therefore, we hypothesized that if the cerebellum contributes to visuo-vestibular interaction, then increased visual dependency would be observed after 1-Hz cerebellar rTMS, as seen in patients with SCA6. We quantified visual dependency when estimating the gravitational vertical before and after rTMS exposure over the posterior cerebellar vermis, which has been shown to be involved in both vestibular (Stephan et al., 2005) and visual motion processing (Cattaneo et al., 2014), and evaluated the effect of rTMS on visual dependency. To confirm the specific involvement of the vermis in the visuo-vestibular interaction, we further examined the effect of rTMS over adjacent cortical areas, namely the early visual cortices (V1–2, Experiment 1) and the cerebellar hemisphere (Experiment 2).

## Materials & Methods

### Participants

In total, 22 (mean ± standard deviation age: 21.3 ± 4.3 years; 10 women, 4 left-handers) and 20 (mean age, 19.8 ± 0.9 years; 3 women, 10 left-handers) healthy volunteers participated in Experiments 1 and 2, respectively. Five participants were included in both the experiments. Handedness was determined using the Edinburgh Handedness Inventory (Oldfield, 1971). All participants reported normal vision and no known neurological, visual, vestibular, or motor disorders (current or historic). The study protocol was approved by the Ethics Committee. Prior to the experiment, all participants provided written informed consent, in accordance with the Declaration of Helsinki.

### General experimental design

In this study, the visual dependency in the perception of gravitational direction was quantified by analyzing the effects of rotating optokinetic stimulation (OKS) on performance in a subjective visual vertical (SVV) judgement task (see below), and the effects of rTMS to specific brain regions on task performance.

Experiments 1 and 2 consisted of three and two sessions, respectively. In each experimental session, 1-Hz rTMS was applied to the cerebellar vermis (vermis condition), early visual cortex (V1– 2 condition), or air (sham condition) in Experiment 1, whereas it was applied to the right cerebellar hemisphere (hemisphere condition), or to the air (sham condition) in Experiment 2. These experimental sessions were conducted on different days, with intervals of at least three days to avoid carry-over effects of prior rTMS. The order of the rTMS conditions was randomized across participants.

On each experimental day, the participants performed the SVV judgement task for 96 trials in the no-OKS condition with a stationary visual background (annulus of dots), followed by 96 trials in the OKS condition with the visual background rotating in the frontal plane (“SVV task session” in Fig. 1A, upper panel). This combination was performed before and after rTMS exposure for 15 min, i.e., 96 trials × 2 OKS conditions × 2 phases = 384 trials for each day (Fig. 1A, upper panel). During the rTMS session, participants were asked to relax in a prone position on the bed. To familiarize themselves with the experimental task, participants performed 14 trials of the SVV judgement task in each condition (No-OKS and OKS conditions), prior to the pre-rTMS phase, on each day (“Task familiarization” in Fig. 1A, upper panel).

**Figure 1.**
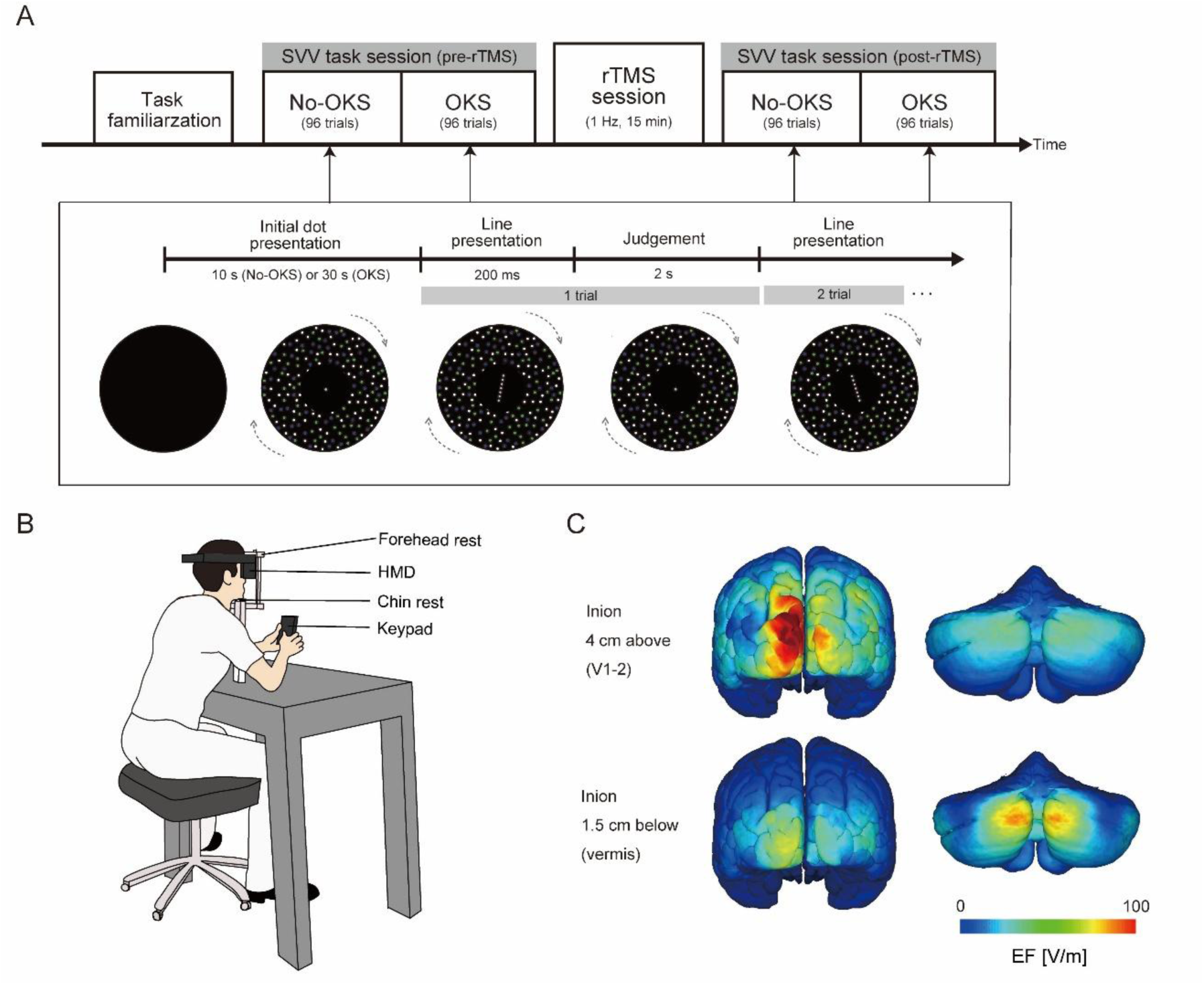
(A) Task procedure on each experimental day. In each SVV task session before or after the rTMS session, the participants judged whether a presented line of dots was tilted in the CW or CCW direction with respect to the gravitational vertical, whereas visual background dots were either rotated (OKS condition for CW group shown as an example; denoted by the grey dashed arrows) or not rotated (No-OKS condition). They performed 96 trials for each SVV task session. One of the three rTMS conditions (vermis, V1–2, sham) was assigned to the rTMS session. (B) Experimental setup. Participants wore the HMD goggle and held the keypad to indicate line orientation during the SVV task session. (C) The estimated EF distribution (V/m) for each coil placement assigned to the V1–2 or vermis condition (see Extended Tables 1-1 and 1-2 for details of the maximum/mean EF in each ROI). Additionally, see Extended Data Figure 1-1 for the EF distribution for the hemisphere condition (1 cm below and 3 cm to the right of the inion) in Experiment 2. EF, electric field; SVV, subjective visual vertical; OKS, optokinetic stimulation; rTMS, repetitive transcranial magnetic stimulation; CW, counter-clockwise; CCW, counter-clockwise; HMD, head-mounted display; ROI, region of interest.

In each experiment, the participants were equally and randomly assigned to either the clockwise (CW) or counter-clockwise (CCW) group. For the CW and CCW groups, CW and CCW OKS was provided in the OKS condition (see *SVV judgement task*) across all phases and rTMS conditions, respectively. This design allowed for a reduction in the number of trials, which helped avoid lengthy experimental times for each participant.

### SVV judgement task

Participants sat on a chair with a head-mounted display (HMD; Oculus Rift S; Meta, Menlo Park, CA, USA), placed their chin on the chin rest, and pressed the superior anterior portion of the HMD against the forehead rest combined with the chin rest during the sessions (Fig. 1B). A non-slip tape was attached between the HMD and forehead rest to minimize involuntary head movements that can be induced by OKS. The head tilt angle in the frontal plane was continuously monitored via the HMD device, and if the head angle was larger than 1° in the CW or CCW direction during the SVV judgement task, the experimenter manually adjusted the participant’s head to an upright position. During the task session, white noise was provided through the HMD to avoid the surrounding auditory cues.

In each SVV task session, the No-OKS condition was initiated by the appearance of an annulus of multicolored (green, white, blue) stationary dots, serving as visual background (outer diameter, 105.5°; inner diameter, 17.9° [viewing angle]; density, 76.9 dots/m^2^), along with a gray fixation dot (diameter, 0.9° [viewing angle]) in the center of display for 10 s (“Initial dot presentation” in Fig. 1A, lower panel). Then, instead of the fixation point, a virtual line consisting of a row of seven gray dots (each dot size, 0.9°; total length of the line, 8.1° [viewing angle]) was presented for 200 ms at a certain tilt angle (“Line presentation” in Fig. 1A, lower panel). Dots were used instead of a solid line to avoid orientation cues that could arise from the visual aliasing of a solid line (Dakin et al., 2018). The tilt angle of the visual line was randomly selected from a set of 12 angles: ±16.5°, ±8.5°, ±4.5°, ±2.5°, ±1.5°, and ±1° relative to the vertical. Positive and negative signs indicate the CW (rightward) and CCW (leftward) directions, respectively. Then, a gray fixation point appeared in the center for 2 s (“Judgement” in Fig. 1A, lower panel), within which the participant was instructed to judge whether the line orientation was tilted in the CW or CCW direction with respect to the gravitational vertical as quickly and accurately as possible by pressing one of two keys on a customized numeric keypad (TK-TCM011SV; Elecom, Osaka, Japan) with the right and left thumbs. Participants were also asked to avoid re-pressing the button during this phase; if they pressed the button, the last button pressed was recorded as the judgement. After the fixation point had disappeared, the visual line was presented again at a certain angle for the next trial. Each line angle was presented twice per block, resulting in 24 trials per block. This block was repeated four times in the No-OKS condition, for a total of 96 trials.

After all trials in the No-OKS condition were completed, the OKS condition was started. Prior to the presentation of the virtual line for the SVV judgement task, the visual background (annulus of dots) was rotated in the CW or CCW direction (depending on the participant’s group) with a center fixation point for 30 s (“Initial dot presentation”). This pre-exposure period provides a stable assessment of the effect of OKS on SVV performance, considering a previous study (Dakin et al., 2018) that showed that the SVV bias toward OKS increased with exposure time but reached a saturation point at 30 s. The angular velocity of the OKS was 30°/s, which could effectively shift the SVV in the direction of the OKS (Dichgans et al., 1972). The SVV trial was then started, with 96 trials completed under the OKS condition and with the same procedure as in the No-OKS condition. Before the SVV task session, the participants were instructed to perform the SVV judgement task without paying attention to the visual background. The total duration of each SVV judgement task session was approximately 10 min.

### TMS

#### Stimulus parameter

In each rTMS session, low-frequency (1-Hz) rTMS was applied to the target cortical site (see below) for 15 min (i.e., 900 pulses) using a MagPro R20 magnetic stimulator (MagVenture, Llandeilo, United Kingdom) with a double-cone coil (D-B-80; MagVenture, Farum, Denmark).

The stimulus intensity of rTMS in each experiment was determined according to each individual’s resting motor threshold (rMT; Cattaneo et al., 2014; Fiori et al., 2015), which was measured on the first day before the experiment. The electromyographic signal of the right first dorsal interosseous muscle was acquired via Ag/AgCl surface electrodes, amplified, bandpass filtered between 15 and 3 kHz with an amplifier (MEG-1200; Nihon Koden, Tokyo, Japan), and sampled at a rate of 10 kHz using a PowerLab 800S A/D converter (ADInstruments, Sydney, Australia). The motor hotspot, i.e., the cortical spot where TMS produced the maximum amplitude of motor-evoked potentials in the right first dorsal interosseous muscle, was explored by delivering single-pulse TMS to various positions around the left primary motor area. The rMT was defined as the minimum intensity that evoked motor-evoked potentials > 50 µV for at least 5 of 10 stimulations applied to the motor hotspot during resting (Rossini et al., 2015). Concerning the coil orientation for rMT determination, the handle pointed backward and laterally at a 45° angle to the nasion–inion line (Mills et al., 1992). The intensity of TMS stimulation was set to 90% of the rMT for each participant. The mean (± standard deviation) intensity was 34.0 ± 5.7% of the maximum stimulator output.

#### Determination of coil placement

In *Experiment 1*, two stimulation sites were targeted: the cerebellar vermis (vermis condition) and the V1–2 regions of the visual cortex (V1–2 condition). To determine the TMS coil positions for these stimulation conditions, we simulated the electric fields (EFs) induced by TMS, using high-resolution head models at the group level.

The head models were constructed with a spatial resolution of 0.5 mm, derived through the segmentation of T1-and T2-weighted structural magnetic resonance images (MRIs) of 18 male participants (21-55 years) were obtained from a freely available repository (NAMIC: Brain Multimodality, 3.0 T MRI scanner, 1 mm voxel size, available online at http://hdl.handle.net/1926/1687). Detailed explanation of head models can be found in our previous study (Laakso et al, 2015). Each segmented head model included detailed anatomical structures, with electrical conductivities assigned to individual tissue types (e.g., gray matter, white matter, cerebrospinal fluid, and bone) based on the established literature values. The development and validation of these models followed the procedures detailed in previous studies (Laakso et al., 2015; Laakso et al., 2018). For volume conductor analysis, the scalar potential finite difference method (Dawson & Stuchly, 1996), with a multigrid algorithm (Laakso & Hirata, 2012), was used to compute the induced EF. The electrical conductivity was assigned to each tissue based on the values provided by Gabriel et al. (1996). This approach calculates the EF distribution by solving the quasi-static Maxwell equations with the vector potential of the TMS coil as the input parameter. A simulation was executed for each head model to predict site-specific EF distributions under the given coil position. As a post-processing step, the EF values from each personalized head model were transformed into a standard brain space, allowing for analysis at the group level. The stimulation sites, the cerebellar vermis and V1–2 regions, were defined anatomically based on standard brain atlases and functional relevance to the experimental objectives (Fischl, 2012).

To determine the coil placement, points were first marked along the inion–nasion line of the head model between 2 cm below and 5 cm above the inion at 0.5-cm intervals. Regions of interest (ROIs) were set in the vermis VII region for the cerebellar vermis, and the combined region of the lingual gyrus (Brodmann 17 area) and cuneus (Brodmann 18 area) for the V1–2 region. The maximum and mean EF values in each ROI were calculated when the TMS coil was placed at each point (Extended Data Table 1-1). To maximize the focality of the stimulation, we selected the point 1.5 cm below the inion to stimulate the vermis, whereas the point 4 cm above the inion was stimulated in the V1–2 condition based on the EF data. We verified that these coil placements ensured a high EF at the target ROIs while keeping the EF at non-target ROIs relatively low (Fig. 1C; see Extended Data Table 1-2).

The coil orientation for stimulation was set as the handle was oriented upward in both the vermis and V1–2 conditions (Cattaneo et al., 2014). In the sham condition, the coil was set at the same position as in the vermis condition, but the coil surface was tilted backward at 90° relative to the scalp surface (i.e., the handle was oriented backward); thus, the brain was not stimulated.

In *Experiment 2,* although the stimulation parameters were adapted from Experiment 1, the target stimulation site was the right cerebellar hemisphere (hemisphere condition). In this condition, the TMS coil was placed 1 cm inferior to the inion on an inion–nasion line and 3 cm lateral to the right, as in previous studies (e.g., Matsugi et al., 2024). EF simulation with the ROI set over the entire right cerebellar hemisphere showed that this coil placement induces a strong EF in the right hemisphere (mean EF, 18.0 V/m; maximum EF, 127.0 V/m), whereas the EFs in the vermis was weaker (mean EF, 26.4 V/m; maximum EF, 54.3 V/m) than that in the vermis condition (mean EF, 34.8 V/m; maximum EF, 68.3 V/m; see Extended Data Fig. 1-1). The coil orientation for this condition was identical to that for the vermis condition. In the sham condition, although the coil position and handle orientation were identical to those in the hemisphere condition, the coil surface was tilted backward at 90° relative to the scalp surface.

### Data Analysis

Responses (CW or CCW) were measured in each trial of the SVV judgement task. If there was no response within the judgment time limit (2 s), the trial was excluded from further analysis.

The proportion of CW responses in each line orientation was calculated for each phase and the rTMS condition. We then estimated the psychometric function by fitting a cumulative Gaussian function to each individual’s response data using the Psignifit 4 toolbox (Schütt et al., 2016) in MATLAB (MathWorks, Natick, MA, USA). Based on the computed psychometric function, we calculated the *SVV* and *judgment precision* values as indices of task performance (see below for details).

#### SVV

The SVV value was determined as the angle at which the visual line was equally likely to be judged as tilted in the CW or CCW direction with respect to the gravitational vertical. Positive and negative values of the SVV indicated deviation in the CW and CCW direction, respectively. Using SVV data, we performed the following steps to quantify the effect of rTMS on visual dependency for each participant. First, the effect of the OKS on the SVV value (*OKS-effect*) was calculated for each phase by subtracting the SVV value in the No-OKS condition from that in the OKS condition. Next, the positive–negative sign of the OKS-effect was reversed only for the participants in the CCW group, so that a positive OKS-effect value indicated a shift of SVV towards the direction of OKS (i.e., a larger OKS-effect value indicates higher visual dependency) for all participants. We confirmed that the OKS-effect was not significantly different between groups in all phases and rTMS conditions in either experiment (Student’s t-test, all *p* > 0.05), indicating that the magnitude of the OKS-effect did not depend on the rotational direction of the OKS (i.e., participant group). Finally, *ΔOKS-effect*, an indicator of the effect of rTMS on visual dependency, was calculated by subtracting the OKS-effect in the pre-rTMS phase from that in the post-rTMS phase for each rTMS condition.

#### Judgement precision

The cerebellum, including the vermis, has been shown to contribute to the processing of vestibular signals (Dahlem et al., 2016; Bronstein et al., 2008; Bertolini et al., 2012; see also Barmack, 2003 for a review), and the cerebellar rTMS may thus influence vestibular processing. The maximum likelihood estimation (MLE) framework assumes that the brain determines the weight of each sensory signal when integrating them according to their reliability, which is the inverse of sensory noise/uncertainty (Angelaki et al., 2009; Ernst & Banks, 2002), i.e. the more reliable the cues, the greater the sensory weight. Based on this framework, if cerebellar rTMS increases/decreases the noise (reliability) of vestibular signals during visual vertical judgement, it may result in an alteration of the visual weight (i.e., visual dependency). A previous study has shown that precision in the estimation of visual vertical in the absence of visual background cues may mainly reflect the noise in the vestibular (otolith) signals (Tarnutzer et al., 2009), as also demonstrated in patients with vestibular loss showing imprecision in visual vertical estimation (Bürgin et al., 2018). Therefore, to assess the effect of rTMS on the vestibular noise underlying the estimation of visual vertical, we quantified the precision in the SVV judgement task by calculating the just noticeable difference (JND; Barnett-Cowan et al., 2010) in the No-OKS condition and evaluated the effect of rTMS.

Based on the fitted psychometric function, the stimulus values (i.e., line angles) at response rates of 25% (*LA*_25_) and 75% (*LA*_75_) were estimated, and the JND in each phase and rTMS condition was then calculated as follows (Gescheider, 1997):

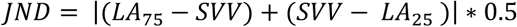

Larger JNDs indicate less precise judgment. As in the SVV task, the effect of rTMS on the JND (*ΔJND*) was computed by subtracting the JND in the pre-rTMS phase from that in the post-rTMS phase.

#### Statistical analysis

To evaluate the group effect of rTMS to the cerebellar vermis or V1–2 on each judgement performance measure (*ΔOKS-effect, ΔJND*), we conducted statistical analysis using a linear mixed-effect model (LMEM) for each performance using the lme4 package (Bates et al., 2015) implemented in R (version 4.3.3; R Development Core Team, Vienna, Austria). The model for Experiment 1 is as follows:

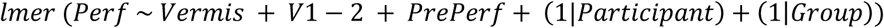

where *Perf* represents each performance metric (ΔOKS-effect or ΔJND) as the dependent variable, and *Vermis* and *V*1 − 2 are binary-coded predictors indicating the rTMS condition. Specifically, the *Vermis* term was coded as 1 when rTMS was applied to the vermis and as 0 otherwise, whereas the *V*1 − 2 term was coded as 1 when rTMS was applied to V1–2 and as 0 otherwise. The sham condition was represented by 0 for both terms, serving as the reference condition. We also included each performance in the pre-rTMS phase (i.e., *OKS-effect_pre_* or *JND_pre_*, denoted in the equation by *PrePerf*) into the model as a fixed effect to control for the effect of baseline performance before rTMS exposure, which may affect the effect of rTMS on performance (Corp et al., 2020). Additionally, two factors, participant and group (CW or CCW), were included as random intercepts (*Participant*, *Group*).

As mentioned above, an LMLE analysis was also performed for ΔOKS-effect in Experiment 2. In this model, two fixed-effect factors, rTMS over the cerebellar hemisphere (*Hemisphere*) and OKS-effect in the pre-rTMS phase (*OKS-effect_pre_*), and two random intercepts (*Participant*, *Group*) were included. The effect size (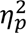) of each factor in the LMEM was calculated. The significance level for all statistical analyses was set at *p* < 0.05.

## Results

### Experiment 1

Data from two participants were excluded from the group analysis because of significant data loss (approximately 1/3 of all trials; ID: 9) and non-completion of the experiment because of nausea (although without seizures or syncope) during OKS exposure (ID: 13). After excluding trials with no response within the time limit (2 s), 99.3% of the total trials were included in the group analysis.

Figure 2A shows the proportion of CW responses at each line angle and the fitted psychometric functions of a representative participant (ID: 3; CW group) for each OKS condition and the rTMS phase in the sham condition. The SVV value was nearly zero in the No-OKS condition for both rTMS phases (pre-rTMS phase, −0.73°; post-rTMS phase, −2.18°), indicating accurate estimation of the visual vertical without visual rotational cues. In the OKS condition, the proportion of responses opposite (i.e., CCW) to the OKS direction increased, leading to SVV shifts toward the OKS (i.e., CW) direction (pre-rTMS phase, 4.19°; post-rTMS phase, 4.04°). This shift was confirmed at the group level by the positive (CW group) and negative SVV angles (CCW group), regardless of the rTMS condition or phase (Table 1).

**Figure 2.**
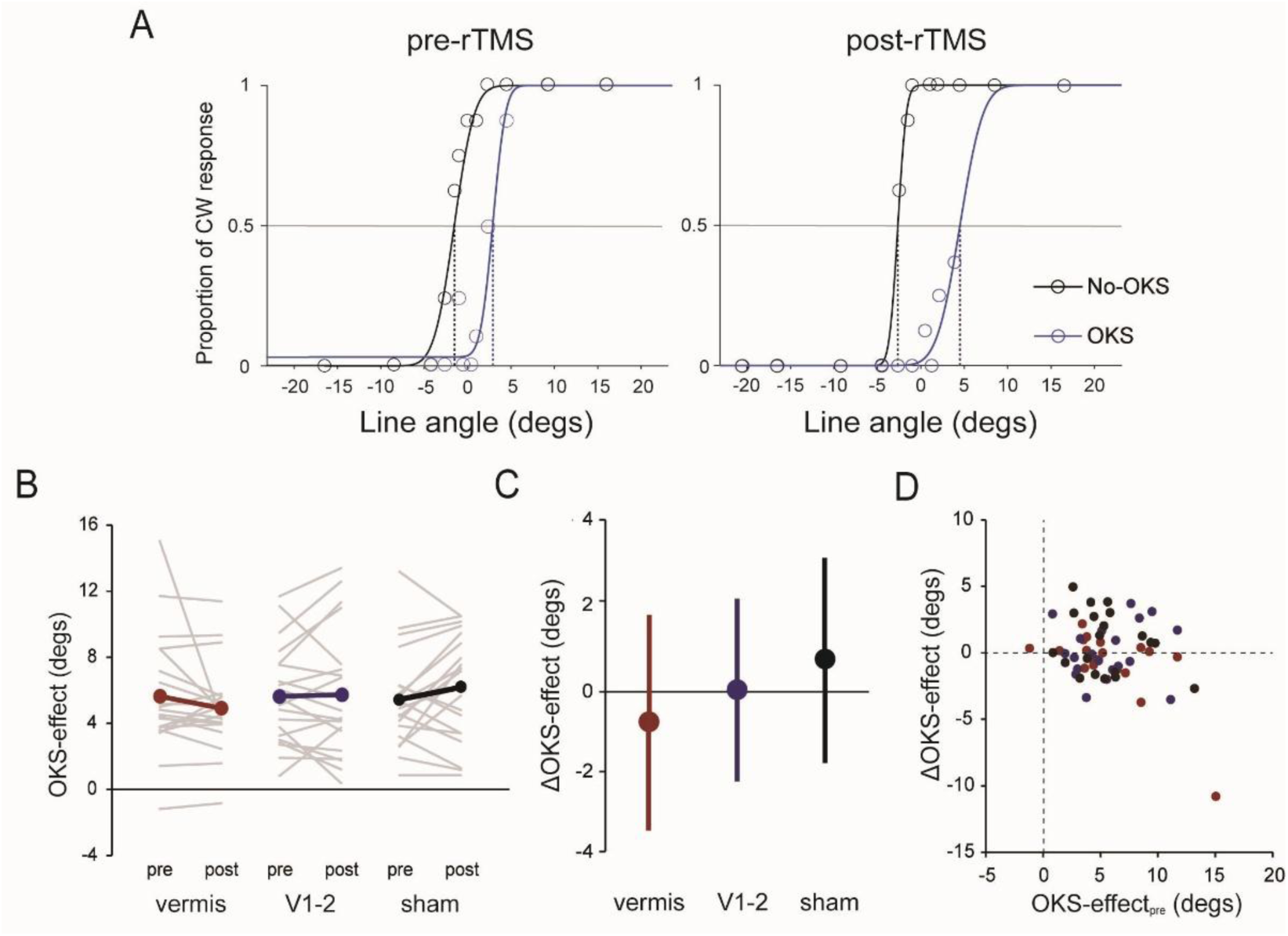
(A) SVV task performance in each OKS condition of the sham condition for a representative participant (Sub ID = 4, CW group). Each color (black, No-OKS condition; blue, OKS condition) of circles and lines represent the proportion of CW responses for each line angle and fitted psychometric function, respectively. The intersection of each psychometric function with the horizontal grey line (response proportion = 0.5) indicates the SVV angle, as represented by the bottom of each dashed line. (B) The OKS-effect on SVV for each phase and rTMS condition. Grey thin lines and colored (red, vermis; blue, V1–2; black, sham) thick lines represent individual and group mean values, respectively. (C) ΔOKS-effect for each rTMS condition. Error bars represent the standard deviation. (D) Relationship between the OKS-effect in the pre-rTMS phase (OKS-effectpre) and ΔOKS-effect. Each plot shows the individual-level data for each rTMS condition (red, vermis; blue, V1–2, black, sham). SVV, subjective visual vertical; OKS, optokinetic stimulation; rTMS, repetitive transcranial magnetic stimulation; CW, clockwise.

**Table 1.** Mean SVV (degrees) of CW and CCW groups for each rTMS condition and phase in Experiment 1.

|  |  | vermis |  | V1 |  | sham |  |
| --- | --- | --- | --- | --- | --- | --- | --- |
|  |  | pre-rTMS | post-rTMS | pre-rTMS | post-rTMS | pre-rTMS | post-rTMS |
| CW group | No-OKS | -1.37<br>(2.52) | -0.71<br>(2.26) | -1.57<br>(1.16) | -0.89<br>(1.93) | -1.33<br>(2.32) | -1.25<br>(3.32) |
|  | OKS | 5.06<br>(5.52) | 4.64<br>(3.17) | 4.36<br>(3.91) | 5.44<br>(5.04) | 4.53<br>(4.94) | 5.09<br>(5.21) |
| CCW group | No-OKS | 0.05<br>(1.76) | 0.19<br>(1.95) | 0.86<br>(1.86) | 1.03<br>(2.11) | -0.24<br>(1.77) | 0.02<br>(1.45) |
|  | OKS | -4.83<br>(3.28) | -4.30<br>(3.90) | -4.47<br>(3.24) | -4.10<br>(4.94) | -5.25<br>(3.74) | -5.99<br>(4.46) |

The values in parentheses represent the standard deviation. CW, clockwise; CCW, counter-clockwise; OKS, optokinetic stimulation; rTMS, repetitive transcranial magnetic stimulation

Although a positive OKS-effect (i.e., SVV shifts to the OKS direction) was observed for all rTMS conditions and phases, the OKS-effect tended to decrease after rTMS exposure in the vermis condition, whereas a slight change or increased OKS-effect were observed in the V1–2 and sham conditions (Fig. 2B), as shown by ΔOKS-effect (vermis, −0.73 ± 2.66°; V1–2, 0.10 ± 2.07°; sham, 0.74 ± 2.30°; Fig. 2C). The LMEM revealed a significant negative effect of *Vermis* (*t* = −2.45, *p* = 0.02, 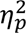 = 0.16), but not of *V*1 − 2 (*t* = −0.10, *p* = 0.33, 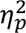 = 0.03), on ΔOKS-effect, independent of the negative coefficient of OKS-effect_pre_ (*t* = −3.82; *p* < 0.001, 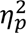 = 0.28; Fig. 2D), implying that ΔOKS-effect was smaller with the larger OKS-effect in the pre-rTMS phase. Thus, visual dependency in the SVV judgement task was attenuated by 1-Hz rTMS over the cerebellar vermis.

Despite the significant effect of vermis rTMS, the ΔOKS-effect in the sham condition was > 0. This result suggests that the significant difference in ΔOKS-effect between the vermis and sham conditions might have been caused by a spurious increase in the OKS-effect in the sham condition rather than the effect of rTMS on the vermis. Considering this possibility, we checked whether a significant change in OKS-effect occurred before and after rTMS exposure in each rTMS condition by conducting additional statistical analysis using the LMEM, with OKS-effect as the dependent variable. As for ΔOKS-effect, *Vermis* and *V*1 − 2 were incorporated as fixed effects and *Participant* and *Group* as random intercepts. In addition, a *Phase* term (coded as 0 for pre-rTMS and 1 for post-rTMS) and interaction terms between *Phase* and *Vermis* or *V*1 − 2 were included as fixed effects. There was a marginal negative interaction between *Phase* and *Vermis* (*t* = −1.94, *p* = 0.055, 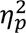 = 0.04) but no significant effects of *Vermis* (*t* = 0.42, *p* = 0.68, 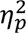 = 0.02), *V*1 − 2 *(t* = 0.37, *p* = 0.71, 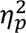 = 0.00), *Phase (t* = 1.38, *p* = 0.17, 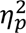 = 0.00), or the *Phase* and *V*1 − 2 interaction *(t* = −0.85, *p* = 0.40, 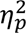 = 0.00). Although this effect was marginal, it suggests that OKS-effect specifically decreased after rTMS was applied to the vermis.

Figure 3A shows the JND results for the No-OKS condition for each rTMS condition and phase. The JNDs tended to slightly increase (i.e., decrease in precision) after rTMS exposure in all rTMS conditions, and the effects of *Vermis* (*t* = −0.32, *p* = 0.74, 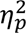 = 0.00) and *V*1 − 2(*t* = 0.47; *p* = 0.64, 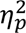 = 0.00) on ΔJND were not significant (Fig. 3B). However, the coefficient of *JND*_*Pre*_ was significant (*t* = −5.13; *p* < 0.001, 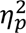 = 0.32), similar to the OKS-effect. These results suggest that precision in visual vertical judgments, mainly based on vestibular inputs without visual rotational cues, may not be affected by rTMS over the cerebellar vermis, in contrast to visual dependency.

**Figure 3.**
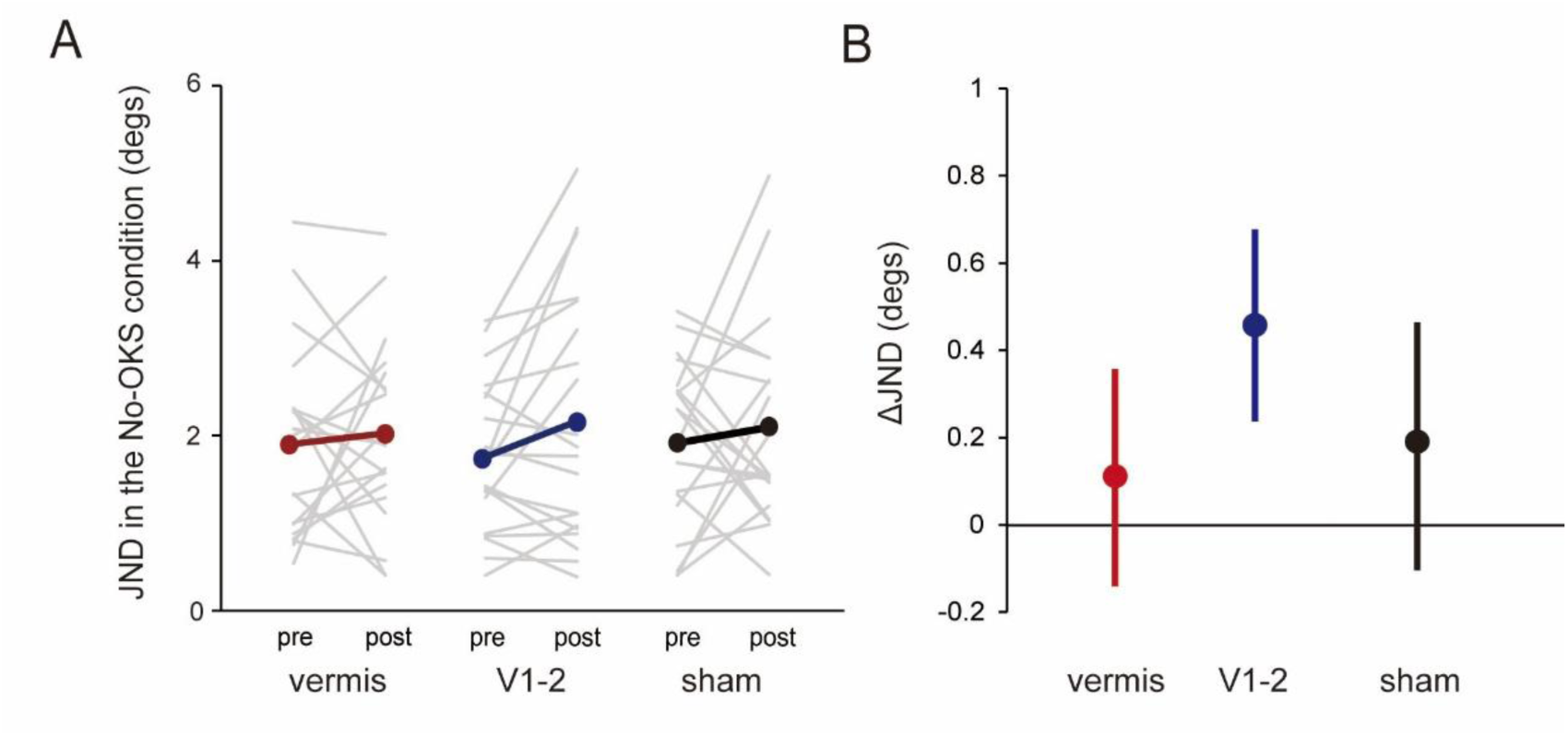
(A) JND in the No-OKS condition for each phase and rTMS condition. Grey thin lines and colored (red, vermis; blue, V1–2; black, sham) thick lines represent individual and group mean values, respectively. (B) ΔJND for each rTMS condition. The error bars represent the standard deviation. JND, just noticeable deference; OKS, optokinetic stimulation; rTMS, repetitive transcranial magnetic stimulation.

We also examined whether rTMS influenced accuracy in the SVV judgement task without visual rotational cues (i.e., SVV in the No-OKS condition). We calculated the change in SVV (ΔSVV) in the No-OKS condition between the phases by subtracting the SVV value in the pre-rTMS phase from that in the post-rTMS phase. ΔSVV was 0.40 ± 1.43° in the vermis condition, 0.43 ± 1.18° in the V1–2 condition, and 0.17 ± 1.39° in the sham condition. LMEM analysis with three fixed effects (*Vermis V*1 − 2, and SVV in the pre-rTMS phase) and two random intercepts (*Participant*, *Group*) showed no significant effect of vermis rTMS (*t* = 1.07, *p* = 0.29, 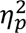 = 0.03) or V1–2 rTMS (*t* = 1.12, *p* = 0.27, 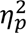 = 0.03), with no significant SVV coefficient in the pre-rTMS phase (*t* = −0.79, *p* = 0.43, 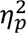 = 0.02). Thus, rTMS to the vermis apparently did not affect the accuracy of the SVV judgement task in the absence of visual rotational cues, consistent with the clinical finding (Tarnutzer et al., 2011) that patients with vestibulo-cerebellar degeneration showed accurate estimation of visual vertical.

### Experiment 2

In total, 99.3% of all the trials were used for the group analysis. Similar to Experiment 1, SVV was nearly zero in the No-OKS condition, whereas SVV was strongly biased toward the direction of OKS in the OKS condition regardless of the rTMS condition or phase (Table 2), supported by the positive OKS-effect (Fig. 4A).

**Figure 4.**
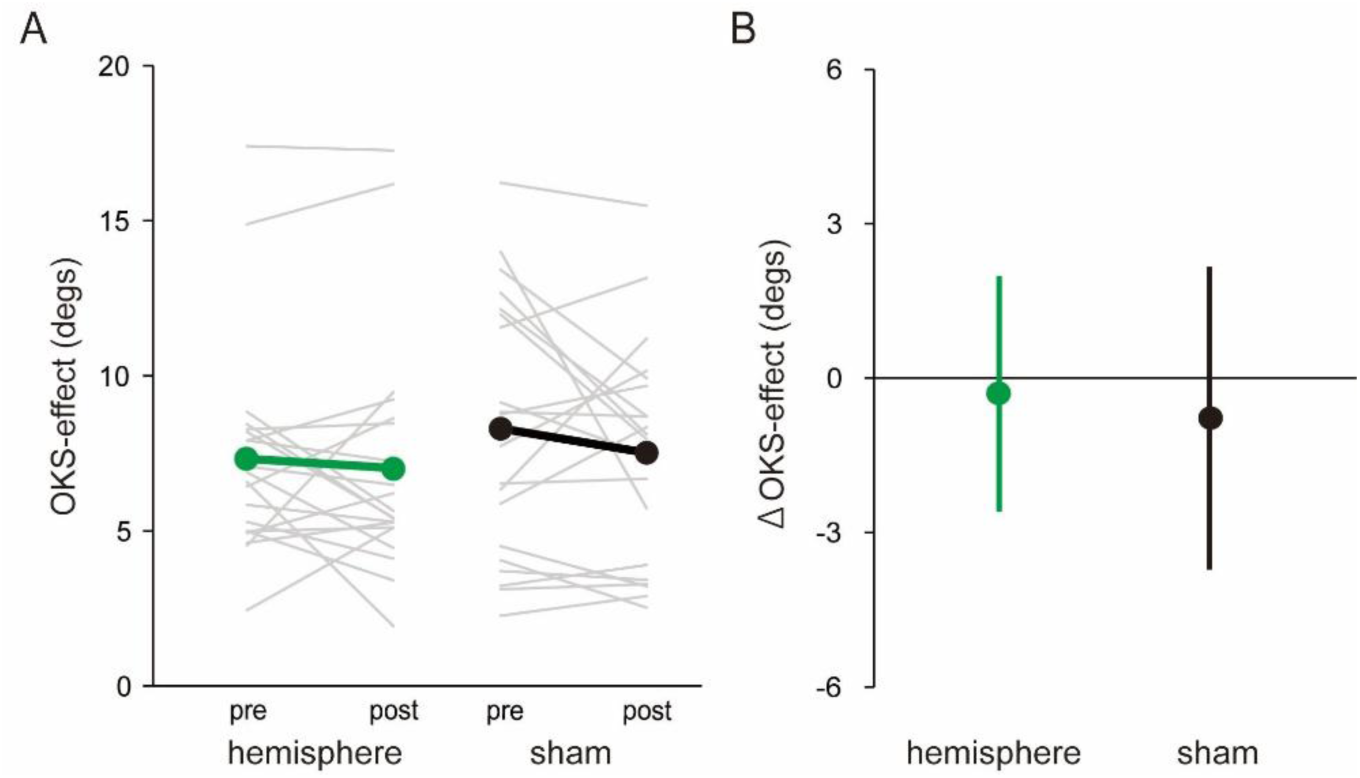
(A) The OKS-effect on SVV judgment for each phase and rTMS condition in Experiment 2. Grey thin lines and colored (green, hemisphere; black, sham) thick lines represent individual and group mean values, respectively. (B) ΔOKS-effect for each rTMS condition. The error bars represent the standard deviation. SVV, subjective visual vertical; OKS, optokinetic stimulation; rTMS, repetitive transcranial magnetic stimulation.

**Table 2.** Mean SVV (degrees) of CW and CCW groups for each rTMS condition and phase in Experiment 2. The values in parentheses represent the standard deviation. CW, clockwise; CCW, counter-clockwise; OKS, optokinetic stimulation; rTMS, repetitive transcranial magnetic stimulation

|  |  | hemisphere |  | sham |  |
| --- | --- | --- | --- | --- | --- |
|  |  | pre-rTMS | post-rTMS | pre-rTMS | post-rTMS |
| CW group | No-OKS | 0.24<br>(1.97) | -0.13<br>(1.55) | 0.56<br>(2.00) | 0.91<br>(3.17) |
|  | OKS | 8.37<br>(4.98) | 7.35<br>(5.43) | 8.62<br>(5.29) | 8.81<br>(5.61) |
| CCW group | No-OKS | 0.42<br>(1.74) | 0.05<br>(2.21) | 0.30<br>(2.53) | -0.40<br>(2.33) |
|  | OKS | -6.09<br>(2.31) | -6.49<br>(2.68) | -8.25<br>(4.27) | -7.55<br>(4.41) |

The mean ΔOKS-effect decreased in the post-rTMS phase for both rTMS conditions (−0.31 ± 2.29° for the hemisphere condition, −0.78 ± 2.94° for the sham condition; Fig. 4B). The LMEM revealed no significant effect of *Hemisphere* (*t* = 0.67, *p* = 0.51, 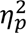 = 0.02) on ΔOKS-effect for the SVV task with no significant coefficient of OKS-effect_pre_ (*t* = −1.56, *p* = 0.13, 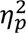 = 0.10). As in Experiment 1, we also checked whether the change in the ΔOKS-effect between the phases was significant for each rTMS condition using another LMEM with two fixed factor (*Hemisphere*, *Phase*), one fixed interaction (*Hemisphere* × *Phase*), and two random factors (*Participant*, *Group*). There were no significant effects of *Hemisphere* (*t* = −1.48, *p* = 0.14, 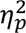 = 0.04) and *Phase* (*t* = −1.17, *p* = 0.25, 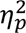 = 0.02) and interaction ( *t* = 0.51, *p* = 0.62, 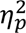 = 0.00). These results indicate that rTMS over the right cerebellar hemisphere did not affect visual dependency for the SVV judgement.

## Discussion

In the present study, to confirm the direct involvement of the cerebellum in the visuo-vestibular interaction for the perception of gravitational vertical, we investigated the effect of 1-Hz rTMS over the posterior cerebellum on visual dependency, quantified by SVV bias induced by rotating OKS. A previous finding that patients with cerebellar degeneration (SCA6) showed exaggerated visual dependency (Dakin et al., 2018), we hypothesized that 1-Hz inhibitory rTMS over the cerebellum could enhance visual dependency in relation to SVV judgement. Contrary to our hypothesis, OKS-induced SVV bias was attenuated after rTMS exposure of the cerebellar vermis. This attenuation was not observed after rTMS over adjacent areas, such as the early visual cortices (Experiment 1) and the cerebellar hemisphere (Experiment 2). Moreover, rTMS to the vermis had no significant effect on judgment precision in the absence of visual motion cues, which is likely to be based on vestibular processing.

Consistent with previous findings (Dichgans et al., 1972; Ward et al., 2017; Niehof et al., 2019; Dakin et al., 2018), the rotating OKS in this study led to a strong SVV bias in the direction of rotation, confirming that the brain integrates vestibular otolith and visual motion cues to estimate gravitational direction (see Dakin et al., 2018 for a review). Importantly, our finding that the OKS-induced SVV bias was attenuated specifically in the vermis condition (Fig. 2) indicates that the cerebellar vermis contributes to visuo-vestibular interactions in relation to the perception of gravitational direction. Animal studies have demonstrated that Purkinje cells in parts of the caudal cerebellar vermis (such as the nodulus and uvula), referred to as the vestibular cerebellum, respond not only to vestibular stimulation, but also to visual motion stimulation (Kano, 1991, 1991; Yakusheva et al., 2013). Human studies have also shown that the cerebellar vermis is activated during visually induced self-motion illusions (Becker-Bense et al., 2012). Furthermore, some neurons in the fastigial nucleus of macaques, which receive inputs from the nodulus/uvula and vermis, were found to respond to OKS (Bryan & Angelaki, 2009).

Dakin et al. (2018) interpreted the increased visual weight (dependency) in patients with SCA as an indirect consequence of the reduced reliability (i.e., increased noise) of vestibular otolith signals, given that an MLE model assuming increased vestibular noise because of a cerebellar deficit, without any effect on visual noise, fitted SVV data from patients with SCA. If inhibitory rTMS to the cerebellum increases vestibular noise, but does not affect visual noise, it should lead to an increase in the visual weight in visual vertical estimation, as observed in patients with SCA. In contrast, OKS-induced SVV bias was attenuated in the vermis condition in this study, leading us to speculate that 1-Hz rTMS to the vermis might have increased the noise in visual motion signals. In fact, Previous studies have shown that the cerebellar vermis is directly involved in processing visual motion signals (Ignashchenkova et al., 2009; Nawrot & Rizzo, 1995, 1998; Cattaneo et al., 2014; Baumann et al., 2015; Bense et al., 2006; Jockisch et al., 2005). Specifically, lesions in the midline cerebellum impair the ability to discriminate the direction of visual motion in humans (Nawrot & Rizzo, 1995, 1998) and monkeys (Ignashchenkova et al., 2009). Additionally, the cerebellar vermis is functionally connected to the visual network (Sang et al., 2012; Kellermnann et al., 2012). In contrast to the OKS-induced SVV bias, we did not observe significant changes in judgment precision (ΔJND; Fig. 3) in the absence of visual motion cues in the vermis condition, implying that vermis rTMS is unlikely to have effectively modulated the vestibular processing underlying visual vertical estimation. Together, these results suggest that 1-Hz rTMS over the vermis might have increased visual noise by interfering with visual motion processing, while having less effect on vestibular processing, ultimately leading to reduced visual weight in visuo-vestibular integration processes.

rTMS over the early visual cortices (V1–2 condition) did not significantly affect visual dependency in this study (Fig. 2C). These regions play an important role in visual motion processing, along with the cerebellar vermis, by projecting local motion signals to extraretinal regions such as the MT/V5 area (Shipp & Zeki, 1989). This is supported by the finding that disrupting V1 by TMS reduces the ability to discriminate the direction of translational visual motion (Laycock et al., 2007; Cattaneo et al., 2014). Given these facts, one may argue that inhibitory rTMS of this region could reduce visual dependency by interfering with visual motion processing, as in the vermis condition. Nevertheless, the lack of a significant effect of the V1–2 rTMS condition is likely attributable to the use of “circular” visual motion. Deutschlander et al. (2004) evaluated brain activity while visually induced self-motion (vection) was perceived about the line of sight (i.e., rotation in the frontal plane) or along the axis (i.e., translation) using positron emission tomography. They found that V1–2 (Brodmann areas 17 and 18) was strongly activated during translational vection, but not during rotational vection, compared to the presentation of stationary dots (i.e., no visual motion cues). Furthermore, Donato et al. (2020) reported that rTMS delivered to V1–2 had no effect on the discrimination of the direction of rotation of a random dot in the frontal plane. These findings suggest that the early visual cortex may not contribute to the processing of circular visual motion, which may be responsible for the lack of a significant effect of V1–2 rTMS on visual dependency in this study.

The effect of rTMS on visual dependency must be discussed carefully, as the difference in the effect of OKS between the pre- and post-rTMS phases in the vermis condition was not significant in this study, although there was a trend toward significance (*p* = 0.054). This lack of significance may be attributed to the difficulty of effectively stimulating the vermis, which is anatomically hidden beneath the cerebellar hemispheres and is located deep in relation to the scalp (Voogd & Glickstein, 1998). In fact, the simulated EF value at the target area was low with coil placement in the vermis condition compared to that in the V1 condition (see Fig. 1C and Extended Data Table 1). This result reflects a limitation of cerebellar TMS in this study; the modulation of neural activity in the vermis may have been limited, potentially resulting in an insufficient effect of rTMS. Given the small effect of vermis rTMS, further studies with a larger sample size may be necessary to confirm the contribution of the vermis to visuo-vestibular interactions.

Our findings do not determine whether the cerebellar vermis plays a direct role in combining and weighing visual and vestibular cues *per se*. Such a direct role may be unlikely, as the MLE model, which assumes changes only in vestibular noise, was found to fit the SVV data of all participants well, regardless of the presence or absence of cerebellar deficits (Dakin et al., 2018). Presumably, other cortical areas may be candidates for visuo-vestibular integration especially for the orientation perception (Cullen, 2019 and Cheng & Gu, 2018 for reviews). Electrophysiological studies in monkeys have demonstrated that neurons in the dorsal region of the medial superior temporal area and ventral intraparietal area are tuned to the direction of self-motion based on both vestibular and visual motion signals (Gu et al., 2006, 2016; Bremmer et al., 2002; Chen et al., 2011). Moreover, recent studies have shown that human homologues of the dorsal region of the medial superior temporal area and ventral intraparietal area may be involved in visual- and/or vestibular-based self-motion perception (Schmitt et al., 2020; Schindler & Bartels, 2018). It is likely that visual and vestibular signals are processed via various cortical and subcortical areas, including the cerebellum, and then integrated into higher-stage cortical areas including these regions to construct an internal representation of the direction of gravity (Kheradmand & Winnik, 2017). Further research is needed to investigate in detail how these cortical areas and the cerebellum interact in the perception of gravitational direction.

In conclusion, the present study showed that visual dependency in visual vertical judgements was attenuated specifically after 1-Hz rTMS over the cerebellar vermis. This finding, although contradicting a previous result in patients with cerebellar degeneration (Dakin et al., 2018), supports the direct involvement of the cerebellum in visuo-vestibular interaction for the perception of gravitational direction.

## Acknowledgments

We thank Dr. Satoshi Tanaka from Hamamatsu University School of Medicine for helpful advice with the design of this study, and Dr. Jose Gomez-Tames from Chiba University for technical assistance. This study was supported by the Japan Society for the Promotion of Science (JSPS) KAKENHI (20K19305 and 23K16761), Otemon Gakuin University (to K.T.), and Shijonawate Gakuen University (to A.M).

