## Supplementary figures and images for "Cerebellum involvement in visuo-vestibular interaction for the perception of gravitational direction: a repetitive transcranial magnetic stimulation study"

### Extended Data Figure 1-1

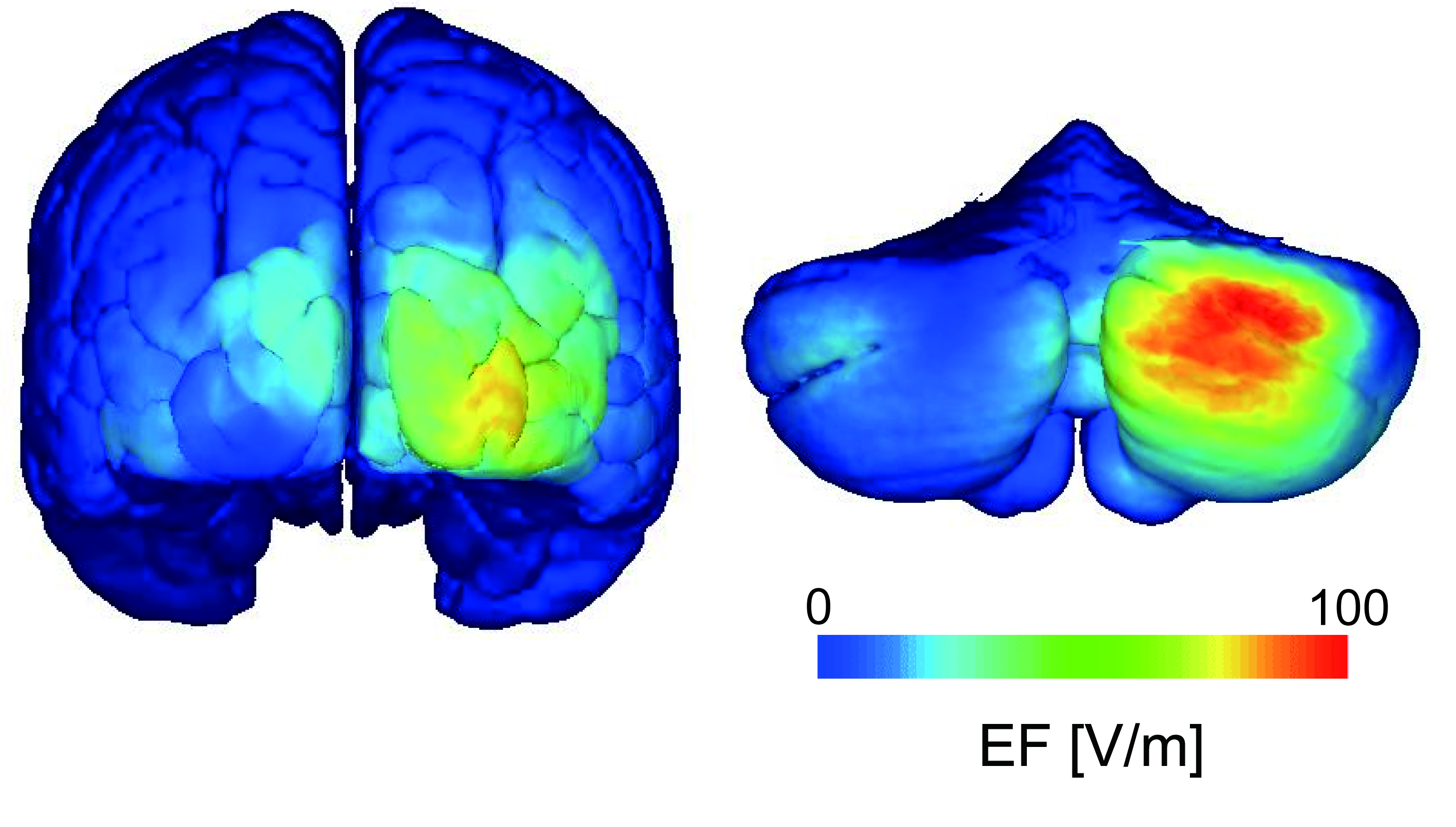
