## Extended Data Table 1-1 for "Cerebellum involvement in visuo-vestibular interaction for the perception of gravitational direction: a repetitive transcranial magnetic stimulation study"

**Table 1-1. Mean and maximum EFs for each point**

|  | V1-2 ROI | | Vermis ROI | |
| --- | --- | --- | --- | --- |
| Distance from the inion　(cm) | Mean  (V/m) | Maximum  (V/m) | Mean  (V/m) | Maximum  (V/m) |
| -2.0 | 33.6 | 72.3 | 34.0 | 67.2 |
| -1.5 (Vermis condition) | 36.5 | 79.4 | 34.8 | 68.3 |
| -1.0 | 39.0 | 85.3 | 35.3 | 68.6 |
| -0.5 | 41.9 | 92.2 | 35.9 | 69.3 |
| 0 | 44.9 | 98.4 | 36.4 | 69.8 |
| 0.5 | 47.2 | 103.3 | 36.4 | 68.9 |
| 1.0 | 49.4 | 107.7 | 36.1 | 67.8 |
| 1.5 | 51.2 | 110.9 | 35.5 | 66.0 |
| 2.0 | 52.2 | 112.4 | 34.5 | 63.5 |
| 2.5 | 52.2 | 111.4 | 32.9 | 59.9 |
| 3.0 | 52.2 | 110.2 | 31.3 | 56.6 |
| 3.5 | 50.8 | 106.1 | 29.4 | 52.6 |
| 4.0 (V1-2 condition) | 49.4 | 101.9 | 27.6 | 49.0 |
| 4.5 | 47.7 | 97.0 | 25.9 | 45.7 |
| 5.0 | 45.4 | 91.3 | 24.2 | 42.4 |
