## Extended Data Table 1-2 for "Cerebellum involvement in visuo-vestibular interaction for the perception of gravitational direction: a repetitive transcranial magnetic stimulation study"

**Table 1-2. Ratio of EF for each point to peak EF**

|  | V1-2 ROI | | Vermis ROI | |
| --- | --- | --- | --- | --- |
| Distance from the inion (cm) | Mean | Maximum | Mean | Maximum |
| -2.0 | 0.643 | 0.643 | 0.934 | 0.963 |
| -1.5 (Vermis condition) | *0.698* | *0.706* | **0.956** | **0.979** |
| -1.0 | 0.747 | 0.759 | 0.970 | 0.983 |
| -0.5 | 0.803 | 0.820 | 0.986 | 0.993 |
| 0 | 0.859 | 0.876 | 1.000 | 1.000* |
| 0.5 | 0.905 | 0.919 | 1.000* | 0.987 |
| 1.0 | 0.947 | 0.958 | 0.992 | 0.971 |
| 1.5 | 0.980 | 0.987 | 0.975 | 0.946 |
| 2.0 | 1.000* | 1.000* | 0.948 | 0.910 |
| 2.5 | 1.000 | 0.991 | 0.904 | 0.858 |
| 3.0 | 0.999 | 0.980 | 0.860 | 0.811 |
| 3.5 | 0.973 | 0.944 | 0.808 | 0.754 |
| 4.0 (V1-2 condition) | **0.945** | **0.906** | *0.758* | *0.702* |
| 4.5 | 0.912 | 0.863 | 0.712 | 0.655 |
| 5.0 | 0.870 | 0.813 | 0.665 | 0.607 |

To check the focality of stimulation for each coil placement (1.5 cm below and 4 cm above the inion for the Vermis and V1-2 conditions, respectively), we evaluated the highest EF among the points (*peak EF*) and then calculated the ratio of the EF at each point to the peak EF for the max and mean EFs and for each ROI (Table 2). The EF ratio was high (above 0.9) for the target ROIs (shown in bold), whereas it was relatively low (below 0.8) for the non-target ROIs (shown in italics) for both conditions. Ratio of EF for the target and non-target ROIs were represented in bold and italics, respectively. *: the point with the peak EF
